# A Potential Method for Identifying Milk Adulteration and Pb(II) Contamination Scenarios Using Principal Component Analysis From Smartphone Photographs

**DOI:** 10.1101/2024.09.16.613186

**Authors:** Alicia Catelyn Chandra, Cheralyn Clarecia Lianto, Felicia Liem Sulimro, Gabriella Anna Santoso, Michelle Aiko Wang, Lie Miah, Norbertus Krisnu Prabowo

## Abstract

Heavy metal contaminants and adulteration in cow milk products are major issues affecting milk safety and quality, posing health risks to consumers of all ages. These contaminants are sometimes difficult to detect with the naked eye and can potentially pass sensory tests, particularly in white cow milk. This research explores the detection of lead(II) poisoning in milk post-production and the adulteration of different milk samples using an alternative approach through chemometric techniques based on RGB and Grey Area image analysis. A controlled photography environment was used. We analyzed over 105 samples of control, adulterated, and lead(II)-added milk in this study using image processing software. Each photograph was analyzed to provide triplicate Regions of Interest (ROI), resulting in a total of 315 statistical datasets. We found that Principal Component Analysis (PCA) effectively clustered control white milk and Pb(II)-contaminated milk. Clusters of different adulterants were recognized simply by feeding RGB and Grey Area data into PCA. However, some clusters, such as mixed chocolate milk and white milk with lead(II) contamination, were not well distinguished. In this early-stage method, a comparison study with infrared spectra will be required in future research. This alternative method shows potential promise for deployment in limited settings for real-world food quality surveillance and regulation.

---

Cow’s milk and dairy products derived from milk are consumed by more than 80 percent of the global population [1], [2]. Well-known contaminants, such as Pb^2+^ and other heavy metals, are commonly found in milk, especially cow milk [3], [4]. Milk adulteration involves the introduction of harmful or foreign substances into milk, including starch [5], detergent [6], rice milk (known as ‘*air tajin*’ in Indonesia) [7], and other milk-like liquids to deceive consumers into buying a product. This “*cheap way*” of extending the shelf life of milk poses potential harm to consumers. Both heavy metal contaminants and adulterant chemicals pose significant health risks [8].

Contaminants may be added to the product intentionally or unintentionally. Despite environmental changes, such as pollution and industrial activities, monitoring the quality and safety of milk remains a continuous challenge. Contaminants may enter the milk supply chain through cattle and contaminated water [9]. The release of heavy metals from the bones of cattle during growth can lead to the presence of toxic elements like Pb^2+^, which can cause serious health issues. Chemicals in adulterants may also cause changes in the milk, reducing its nutritional value by depleting essential proteins, minerals, and vitamins. Consuming contaminated and adulterated milk can result in serious health problems, including gastrointestinal stress, allergic reactions, or long-term issues such as kidney diseases [4].

To address this issue, chemometric techniques have been implemented to create pattern recognition systems in food science and technology, aiding in quality control and food production assessment. For instance, chemometrics is used in the authentication of halal foods [10] and the detection of heavy metal contamination in food production [11]. The use of smartphones in food science and technology is increasing [12]. Previously, image processing software like ImageJ was commonly used in smartphone spectroscopy research for colored samples [13].

However, the integration of smartphone photographs for analytical purposes remains underdeveloped. There is a research gap in practically applying these gadgets to identify specific types of adulteration and contamination. To our knowledge, no studies have explored using direct measurements of RGB values and gray areas as inputs into PCA for classification and identification in the white milk matrix. This study aims to address this gap by exploring an innovative strategy for detecting milk adulteration and Pb^2+^ contamination. This method involves capturing images of milk samples using a smartphone camera and analyzing them with image processing software, specifically ImageJ. The RGB (Red, Green, Blue) values and grayscale intensity values of the images are subjected to exploratory data analysis using Principal Component Analysis (PCA). This study seeks to reveal patterns and relationships within the photographic data that could indicate the presence of adulterants or contaminants. This method has the potential to enhance the sensitivity and reliability of smartphone-based systems, providing a cost-effective approach to analyzing food safety and quality.

This research is focused on the application of chemometrics for detecting and modeling adulteration and Pb(II)-contamination in cow milk using image analysis from smartphone photography approach. Research Question:

- RQ1: To what extent can variations in RGB brightness values and gray areas be attributed to principal component analysis (PCA) when analyzing milk adulteration and Pb^2+^ contamination scenarios in a white milk matrix?
- RQ2: How do RGB brightness values and gray areas differ across various levels of Pb^2+^ contamination in a white milk matrix?

## LITERATURE REVIEW

Milk adulteration and lead (Pb^2+^) contamination pose significant health risks with potentially severe public health effects. Lead contamination in milk is a well-documented neurotoxin with profound effects on neurological development [14]. Lead(II) is not a nutrient and its presence in food is unacceptable. Chronic exposure, particularly in children, is associated with cognitive deficits, developmental delays, and behavioral problems, which can have lasting impacts on IQ and attention span [15]. In adults, prolonged lead exposure is linked to cardiovascular diseases, including hypertension, and renal damage, potentially leading to chronic kidney disease. Lead can cross the placental barrier, leading to adverse pregnancy outcomes such as preterm birth and developmental delays in infants. Furthermore, lead accumulates in the body over time, primarily in bones and teeth, resulting in long-term health consequences as lead is gradually released back into the bloodstream [16].

Various methods have been employed to detect milk adulteration and lead (Pb^2+^) contamination, each with its own advantages and limitations. For detecting milk adulteration, methods such as chemical tests, spectroscopic techniques, and chromatographic analyses are commonly used. Spectroscopic techniques, including Atomic Absorption Spectroscopy (AAS) [17], infrared spectroscopy [18], and UV/Vis spectroscopy [19], are highly sensitive and accurate for detecting heavy metals like lead. However, these methods require sophisticated equipment, are expensive, and may not be accessible in all settings. Chromatographic methods, such as High-Performance Liquid Chromatography (HPLC) [20], are effective for identifying various adulterants and chemical contaminants but involve complex sample preparation and high operational costs.

Exploratory Data Analysis (EDA) involves a range of techniques designed to summarize and explore the underlying structure of data before applying more formal statistical methods [21]. Common EDA techniques include descriptive statistics, such as histograms, scatter plots, and box plots, which help reveal patterns, and trends. Dimensionality reduction techniques, like Principal Component Analysis (PCA), simplify complex datasets by focusing on key components that capture the most significant variance, making it easier to detect patterns and contributing factors [22].

In relation to photography, an image is defined as a grid of square pixels (picture elements) arranged along the x-axis and y-axis [23]. In an 8-bit format, the pixels are filled with intensity values ranging from 0 (black) to 255 (white). In image analysis, these are referred to as gray areas, and the intensity reflects the brightness of black and white pixels [24]. In a true-color image, the pixels are filled with red, green, and blue color intensities, known as RGB values.

## EXPERIMENTAL DETAILS

### General Protocol

The overall research pathway used in the experiment is illustrated in Figure 1. The general protocol for the experiments was followed: all sample preparations and experiments were completed within the same day. Clean volumetric apparatus were used to measure sample volumes. In all variation experiments, the total volume of the liquid was kept constant at 50.0 cm^3^ ± 0.05 cm^3^. During the experiments, all containers were tightly sealed with caps, and any remaining samples were stored in a refrigerator at 4°C. Sterile transparent plastic containers with secure caps (maximum volume 80.0 cm^3^) were used. Only one smartphone with fixed settings was used for the entire image acquisition.

**Figure 1.**
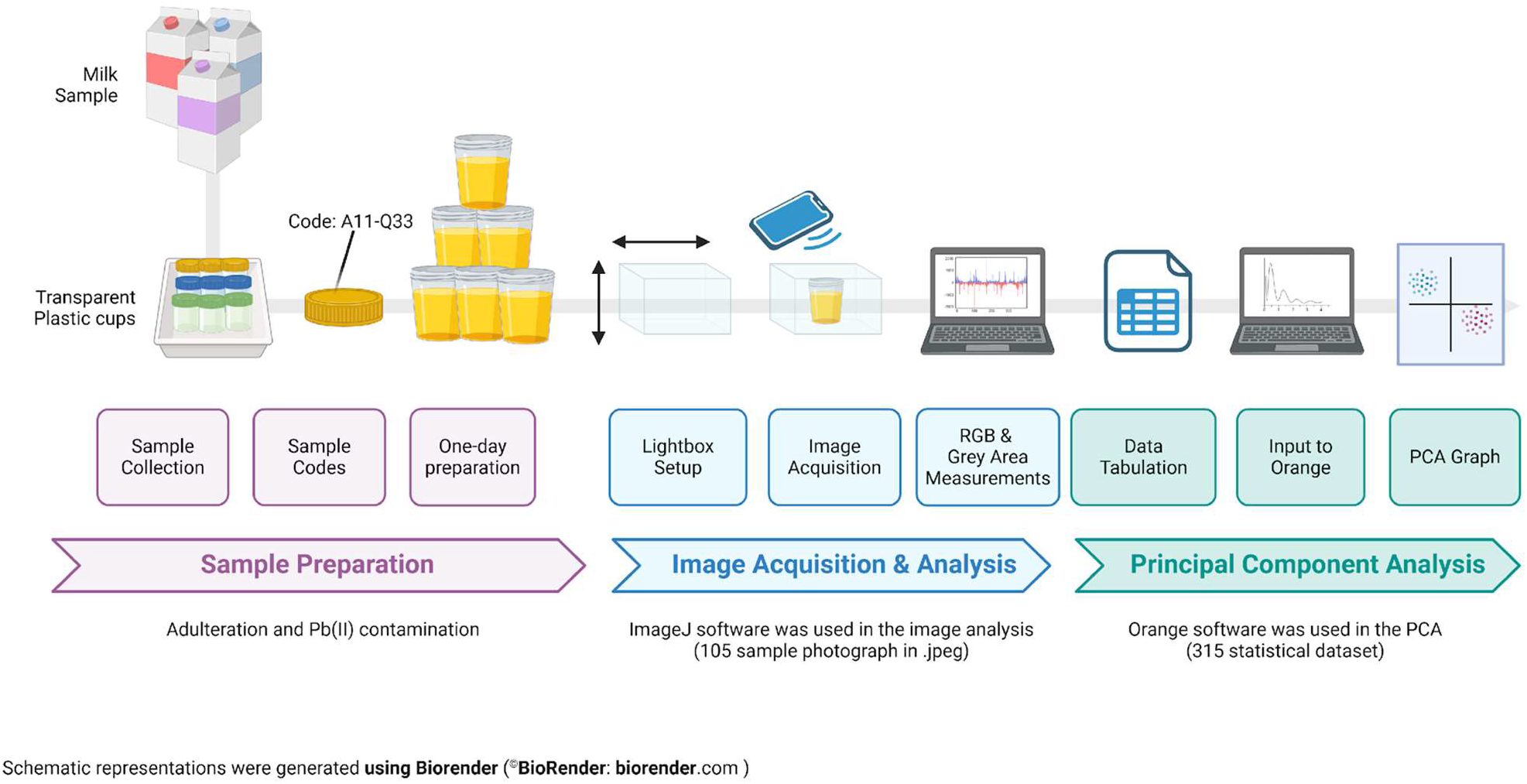
The overall research pathway used in this experiment, with all steps conducted on the same day.

Two main software applications were used in this study: ImageJ for RGB and gray area measurements, and Orange for conducting PCA. ImageJ (from National Institutes of Health) is a widely used open-source software for image analysis [25], [26]. It offers a range of tools for processing and analyzing images, including features for image segmentation, quantification, and enhancement. Orange is an open-source data mining and exploratory statistics software [27].

### Milk Samples and Preparation

According to the literature, processed milk often contains higher levels of contaminants, particularly heavy metals, compared to raw milk [28]. Therefore, we selected commercial processed cow milk products as our samples. The studied matrix was one commercial brand, Ultra Milk^®^ produced by P.T. Ultrajaya Milk Industry & Trading Company, Tbk. This brand was selected in a random lottery from a total of 12 cow milk brands available at convenience stores in the Kelapa Gading area of North Jakarta (July 2024). All of these brands offer various flavors and colorings in their dairy products.

In this study, four different liquid milk products from Ultra Milk^®^ were used as the matrix for the experiments: Ultra Milk Full Cream®, Ultra Milk Taro Flavor^®^, Ultra Milk Strawberry Flavor^®^, and Ultra Milk Chocolate Flavor^®^. The packaging of each product was checked to ensure it was intact, undamaged, and unopened before use. All liquid milk products were used as is, without dilution with distilled or tap water.

For preparation, the four milk products were placed into separate 50-cm^3^ burettes, which were labeled as *Full Cream, Taro, Strawberry*, and *Chocolate*. The *Full Cream* milk was mixed with the other flavors in a 50:50 ratio, resulting in a total volume of 50.0 cm^3^, and was transferred into sterile transparent plastic cups with caps. Different clean glass rods were used for each stirring process to avoid contamination. In total, 17 different sample groups were used in this study, coded from A to Q. The sample coding was straightforward; for example, A13 indicates “A” for control white milk, “1” for the first replication, and “3” for the third photograph. Each identity was sampled at least in triplicate. The sample coding and their identities are tabulated in Table 1.

**Table 1.** Sample Coding and Identity.

| Sample Code | Sample Identity |
| --- | --- |
| A11-A63 | Full Cream Milk<sup>®</sup> (Control) |
| B11-I33 | Various concentrations of lead(II) in Full Cream<sup>®</sup> Milk matrix |
| J11-J33 | Full Cream Milk<sup>®</sup> + Chocolate Flavored Milk<sup>®</sup> (50:50 ratio) |
| K11-K33 | Full Cream<sup>®</sup> + Strawberry Flavored Milk<sup>®</sup> (50:50 ratio) |
| L11-L33 | Full Cream<sup>®</sup> + Taro Flavored Milk<sup>®</sup> (50:50 ratio) |
| M11-M63 | Rice water (*Air Tajin*) (Control) |
| N11-N33 | Full Cream<sup>®</sup> + Rice water (*Air Tajin*) (50:50 ratio) |
| O11-O33 | Chocolate Flavored Milk<sup>®</sup> (Control) |
| P11-P33 | Strawberry Flavored Milk<sup>®</sup> (Control) |
| Q11-Q33 | Taro Flavored Milk<sup>®</sup> (Control) |

Rice water (*air tajin*) is a common adulterant for white milk in Indonesia and is even used as a traditional milk substitute. In this study, rice water was prepared fresh by filtering the colloidal liquid obtained from half-cooked white rice (*Oryza sativa*). White rice was purchased from a traditional market in the Kelapa Gading area of North Jakarta, Indonesia. Four hundred grams of white rice were added to a saucepan with 1500 cm^3^ of distilled water. The mixture was boiled for about 3-5 minutes until the rice was half-cooked and bubbles began to rise. The water turned into a white colloidal liquid, similar in appearance to milk. The mixture was then stirred and filtered to produce rice water (*air tajin*).

Rice water (*air tajin*) was used as an adulterant in Full Cream^®^ samples, with these samples coded N11-N33. The control samples for rice water were coded M11-M63. The sample codes and their tabulation are provided in Table 1, with a fixed total volume of 50.0 cm^3^.

To simulate Pb^2+^ contamination in a white milk matrix, lead(II)-contaminated milk samples were prepared. Solid Pb(NO_3_)_2_ was acquired from Merck and dissolved in 1.00 dm^3^ of Full Cream^®^. This created a stock solution containing 1000 ppm of Pb^2+^. From this stock solution, dilutions were performed in the Full Cream matrix to achieve concentrations of 200 ppm, 300 ppm, 400 ppm, 500 ppm, 600 ppm, 700 ppm, and 800 ppm of lead(II)-contaminated milk. All experimental waste involving lead(II) compounds was collected in a separate container for proper chemical disposal. All laboratory disposal procedures were strictly followed in this study to ensure zero harm to the environment during the experiments.

### Image Acquisition and Analysis

The image acquisition in this study was controlled in an attempt to ensure accuracy, reliability, and reproducibility. A light box, commonly used in photography, was positioned away from direct sunlight and equipped with white LED lights. The dimensions of the light box were 20 cm (length) x 20 cm (width) x 20 cm (height). The LED lights were powered by an AC adapter. The light box was chosen because it has a hole on the top, allowing photographs to be taken from above and from the front side (see Figure. 2).

**Figure 2.**
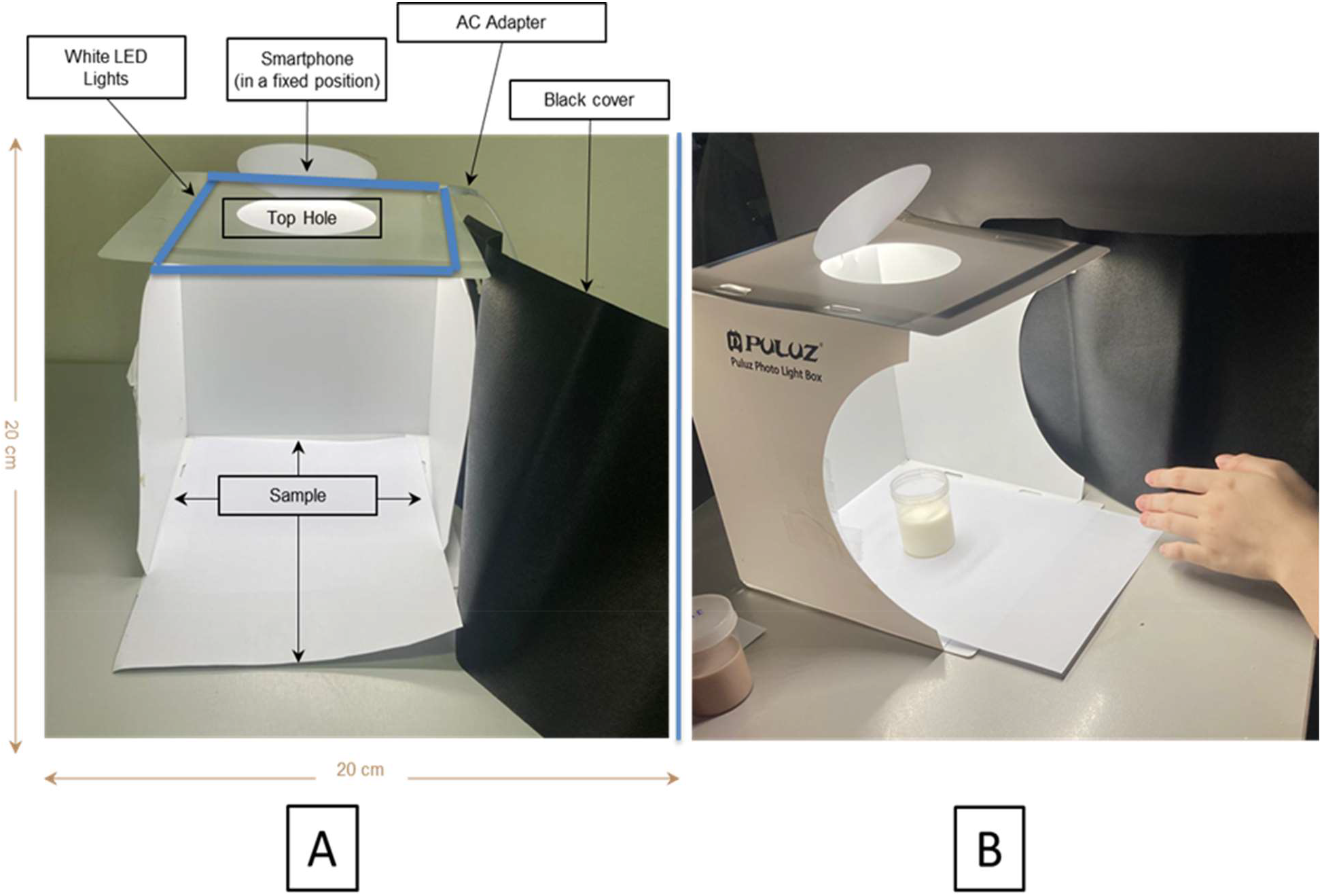
The image acquisition setup. (A). The dimension of the light box used. (B). The sample container was put in the middle of the box (below the top hole). The distance of the smartphone camera and the sample was maintained for each image acquisition.

One smartphone was used for the entire image acquisition process (an iPhone 13 Pro with fixed zoom and brightness settings). The distance of the sample container from the camera was kept constant and regularly checked using a ruler. The distance from the surface of the milk sample to the camera was also kept constant. To avoid exposure from other light sources, the laboratory lights were turned off, and the front of the light box was covered with black cloth to prevent any disturbance in the reflectance color, other than from the sample (see Figure. 2). The obtained images were thoroughly uploaded to Google Drive, including image names for sample coding (see Appendix). A total of over 100 digital photographs (.HEIC format) were used in this study, which were later converted to .jpeg. The RGB and gray area data from the images were extracted using ImageJ software. In each photograph, three random areas in the middle of the cup were selected as regions of interest (ROI) and recorded in a Google Spreadsheet. The ROI size was set to 64 × 64 pixels (width x height), following common quantification practices using ImageJ [29]. The depth of the liquid sample in each transparent plastic container was monitored during image acquisition by placing a ruler in the photo with the sample in the same frame. ImageJ was used to measure the depth of the milk in the container to three decimal places.

### Principal Component Analysis (PCA)

Two spreadsheets were prepared for PCA using Orange software. The first spreadsheet contains all RGB values and gray areas for the 105 samples (from A11 to Q33), aimed at answering RQ1. The second spreadsheet was prepared to address RQ2, containing data on the variation in lead(II) concentration (in ppm). The matrix was *Full cream* milk sample. The detailed procedures are given in the Supporting Information, *vide infra*.

## HAZARDS

Any chemical waste, including milk waste and/or lead (II)-containing waste, must be segregated for proper disposal. Electronic devices, such as smartphones and laptops, should be kept away from chemicals and potential splashes.

## RESULTS AND DISCUSSION

Smartphone colorimetry has been extensively used with transparent solutions containing colored samples [30]. In this study, the cow milk was used as the matrix. Milk is a colloidal mixture, which poses a unique challenge during image acquisition due to its opaque nature. However, colloidal mixtures exhibit the Tyndall effect, where light scattering correlates with reflectance. The reflectance intensity is related to image parameters such as red, green, and blue values (RGB), as well as brightness in the gray areas [31]. The data analysis obtained using ImageJ included R-value, G-value, B-value, R/G, R/B, B/G, and Gray Areas, along with their standard deviations (see Appendix). The standard deviations for the control white milk (A11-A63) were 0.220 for the R-value, 0.203 for the G-value, and 0.195 for the B-value.

The PCA facilitated data reduction from the image parameters collected experimentally, allowing us to better visualize the relationships between variables. In this study, the first PCA was conducted to address RQ1 (see Figure 3). Across all samples, including controls, the first principal components (PC1, PC2, and PC3) explained 92% of the data variability. The plot between PC2 (x-axis) and PC3 (y-axis) in Figure 3 effectively distinguishes the different clusters of samples. Samples with similar color composition and brightness are positioned closer to each other, while those with different chemical compositions produce distinct color compositions and brightness levels.

**Figure 3.**
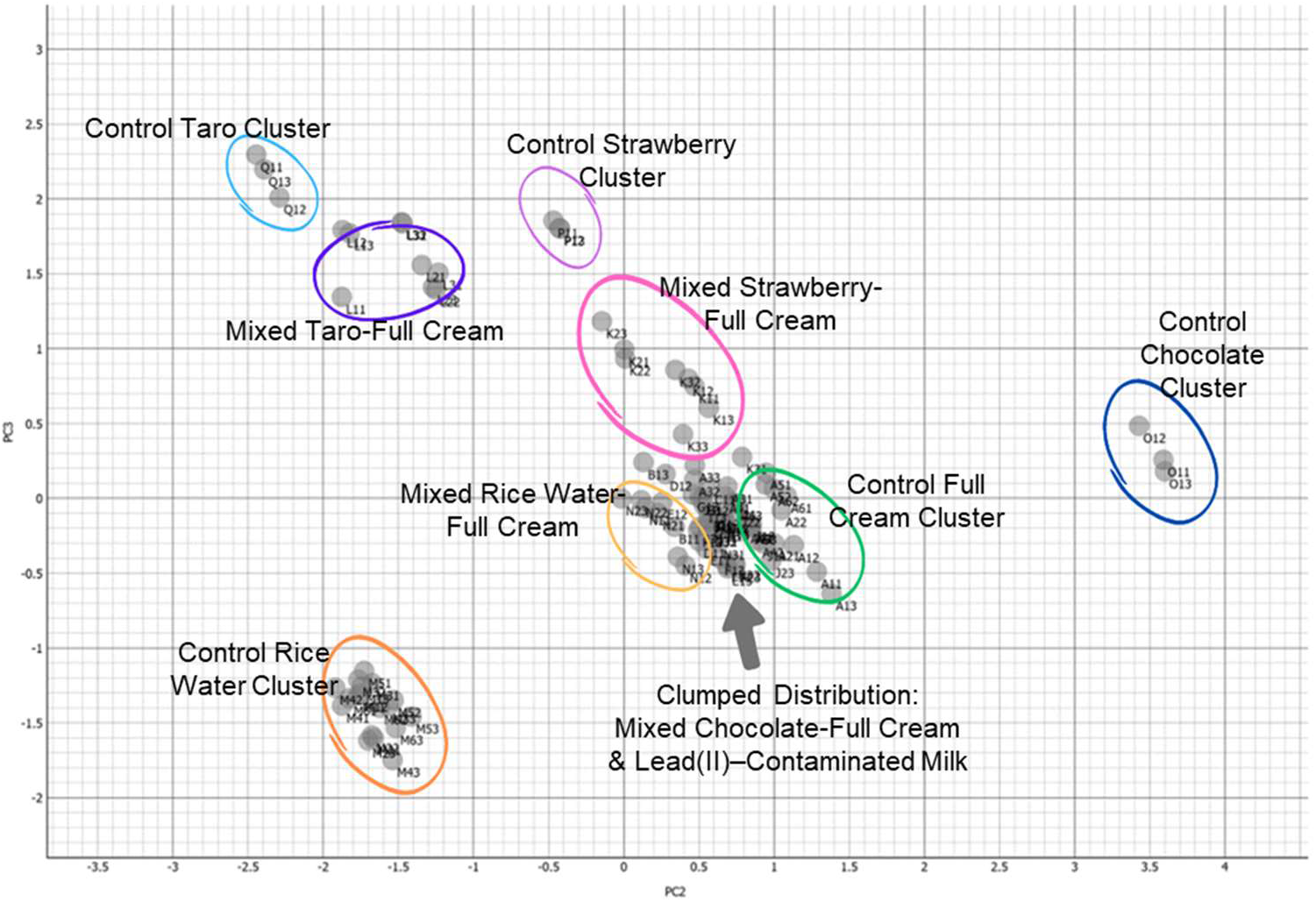
PCA results for the overall samples based on direct input of RGB and grey area values.

In Figure 3, the major contributor to PC2 was the Red/Blue ratio, while PC3 was associated with the Red/Green ratio. The plot shows the data clusters for mixtures of distinct milk flavors—namely full cream, strawberry, chocolate, and taro—along with rice water (*air tajin*) and their respective variations (adulterations). The data cluster for the full cream-strawberry milk mixture was positioned centrally between the clusters for control full cream and control strawberry milk. This observation indicates significant differences in the color and brightness aspects within the groups, enabling us to identify which milk flavors were mixed.

From a quality control perspective, this suggests that clustering helps determine whether milk production is consistent. Consistent production will yield a similar chemical and color composition, resulting in samples clustering closely together. For instance, the mixed rice water-full cream cluster was positioned further from both the control rice water and control full cream clusters. This implies that the incorporation of rice water into full cream milk had a significant impact on the PCA outcomes for the full cream milk. The mixed taro-full cream cluster exhibited a similar pattern to the mixed rice water-full cream cluster, differing in its proximity to the control full cream cluster. Unlike the mixed rice water-full cream cluster, the mixed taro-full cream cluster was positioned closer to the control taro cluster. Despite its equal composition of taro and full cream milk, it displayed results more similar to the control taro milk.

Samples containing a mixture of chocolate and full cream milk exhibited aggregation patterns closest to those of lead(II)-contaminated milk. The Principal Component Analysis (PCA) results for these clusters showed minimal variation, leading to a clumped group distribution. As a result, distinguishing between the milk samples proved challenging due to the minimal differences among them. This suggests that the RGB and gray area measurements were similar between the two groups. However, we found that variations in Pb^2+^ concentration (in ppm) in the milk influenced the Blue/Green ratio. In future studies, a colored background sheet (blue or green) could be placed behind the sample containers (cuvettes or transparent plastic cups) to create discrepancies in the B/G measurements. We speculate that this approach may help better separate the clusters. Additionally, the cluster for the mixed chocolate-full cream milk was notably farther from the control chocolate milk than from the control full cream milk.

The second PCA was conducted to address RQ2, with the results presented in Figure 4. This analysis represents different milk samples intentionally contaminated with Pb^2+^ at various concentrations (in ppm). We aimed to observe how the RGB and gray area measurements responded in the PCA. The investigation through Principal Component Analysis (PCA) effectively clustered and distinctly separated the control (uncontaminated) milk samples (full cream control) from the lead(II)-contaminated milk samples, with 92% of the variability explained. The confidence ellipse in Figure 4 suggests a significant difference in color composition and brightness between the two clusters.

**Figure 4.**
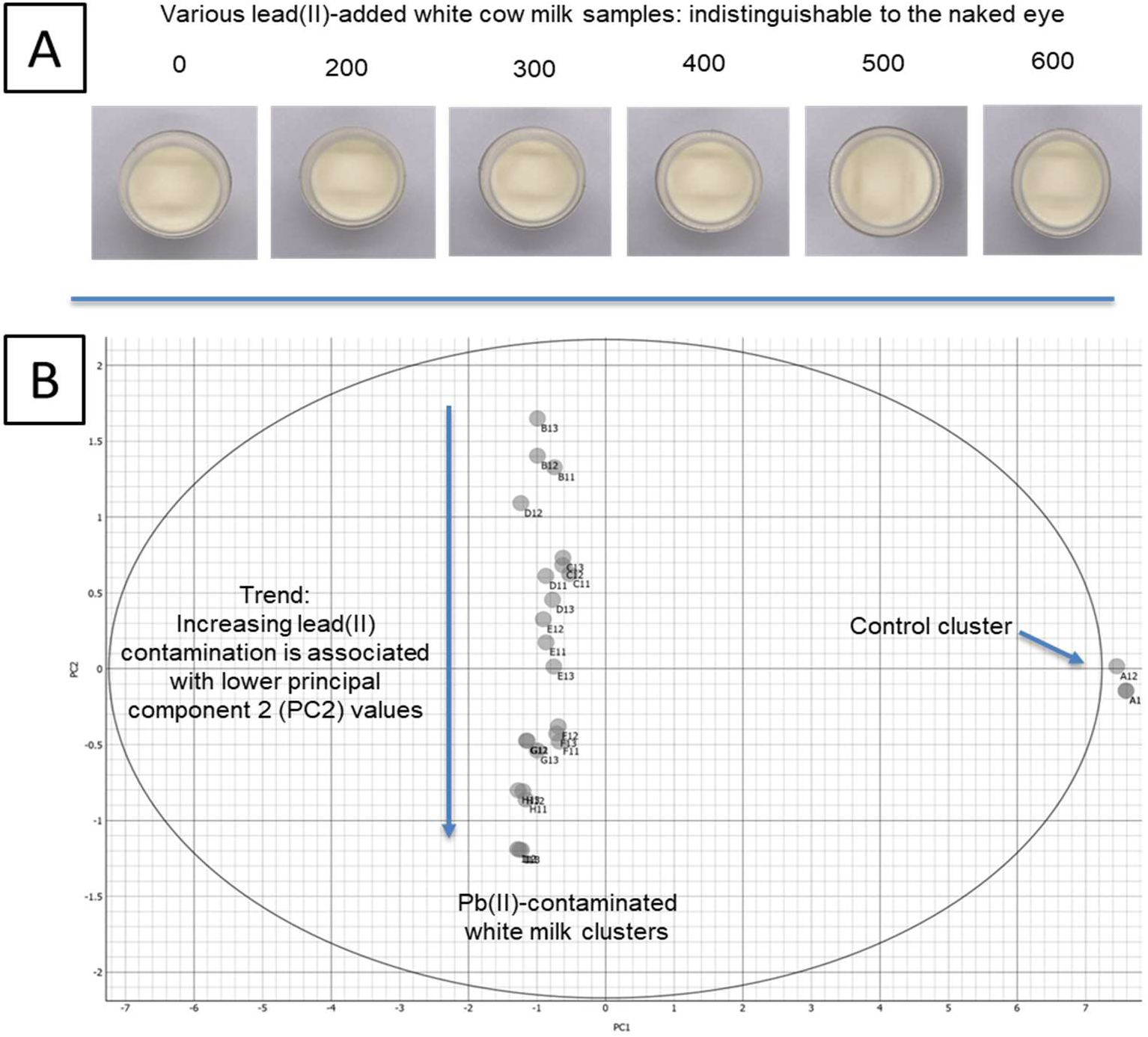
Results for Control and Pb(II)-Added White Milk. (A) Difficulty in comparing the appearance of pure milk samples with lead(II)-added milk samples. (B) Distinct clusters for control and lead(II)-contaminated white cow milk samples.

Additionally, the PCA results show a clear trend: as the Pb^2+^ concentration increases, the PC2 (second principal component) values decrease. However, PC1 does not distinguish between different concentrations of Pb^2+^. For example, 200-ppm lead-contaminated milk appears to have the same color and brightness composition as 500-ppm lead-contaminated milk. Good clustering has been observed in studies analyzing molecular vibrations in lead(II)-contaminated milk products using FTIR-ATR [32]. In milk, Pb^2+^ forms PbCl_2_ and may also create dative bonds (complex ions) with hydroxyl, carboxyl, and amide groups from the protein chains. According to the literature, the presence of Pb^2+^ in milk causes discrepancies in the stretching and bending of hydroxyl groups at 3800–3000 cm^−1^ and the C=O of carboxyl/amide groups at 1700–1400 cm^−1^ [32]. These interactions can only be detected at the molecular level.

This study demonstrates that simple RGB and gray area measurements combined with PCA can effectively sort and cluster various adulterations and lead(II) ion contaminations in milk. This approach could be beneficial for immediate quality control during or after cow milk production. Additionally, analyzing color and brightness characteristics could help ensure the consistency of food products or recipes. In the long run, this method could complement sensory food tests, enabling a more comprehensive assessment of food quality and safety.

## CONCLUSION

Nowadays, smartphones are equipped with cameras and are accessible to almost everyone, offering a commercial opportunity to conduct analytical procedures for managing food quality and safety. In this study, RGB values and gray areas were directly input into PCA to identify adulteration and Pb(II) contamination in white milk. The colorless Pb^2+^ solution, the white precipitate of PbCl_2_, and milk-like adulterants (e.g., rice milk, known as *air tajin*) are difficult to detect visually when mixed with a white milk matrix. However, through a chemometric approach, we were able to differentiate the adulterated milk samples from their controls and distinguish the Pb(II)-contaminated milk from its controls. Notably, the *chocolate-full cream* mixture and Pb(II)-contaminated milk samples clustered together, indicating that they have similar compositions in terms of RGB values and gray areas.

During our research, we encountered challenges in minimizing the impact of external light sources on the RGB values of our samples. Additionally, ensuring that all photos of the samples taken with our smartphones maintained consistent angular precision and dimensions was difficult. For future research, we recommend using a tripod to maintain a fixed distance during image acquisition and working in a more controlled environment. Further research could be enhanced by incorporating standard chemometric analysis with infrared spectra measurements and determining the lower detection limit of the developed method. Several recommendations derived from this research include potential applications in recipe consistency testing, evaluating food appearance and texture quality, assisting with qualitative biochemical identification tests, complementing sensory testing procedures, and identifying additive colorings. A simple smartphone application could provide significant support in this area of research.

## ASSOCIATED CONTENT

## Appendix

- Procedures and tabulation for PCA: https://doi.org/10.5281/zenodo.13766649
- All photographs of samples: https://drive.google.com/drive/folders/1FPigb_tgkGCUseK7qzTy5KT_u2u4RSmH?usp=sharing

## AUTHOR INFORMATION

### Notes

The authors declare no competing financial interest.

## ACKNOWLEDGMENTS

The authors express their gratitude to Prof. Dr. Erdawati, M.Sc., and Dr. Maria Paristiowati, M.Si. from the STEM Education Center at Jakarta State University for their expert guidance on smartphone-based spectroscopy. We also extend our thanks to Ms. Vita Kusumastuti, Mr. Bobby Aprianto, and Mr. David Tan from Badan Pendidikan Kristen PENABUR Jakarta for their unwavering support.

